# The “Alu-ome” shapes the epigenetic environment of regulatory elements controlling cellular defense

**DOI:** 10.1101/2021.11.21.469436

**Authors:** Mickael Costallat, Eric Batsché, Christophe Rachez, Christian Muchardt

**Affiliations:** Institut de Biologie Paris-Seine (IBPS), CNRS UMR 8256, Biological Adaptation and Ageing, Sorbonne Universités, 75005, Paris, France

## Abstract

Promoters and enhancers are sites of transcription initiation (TSSs) and carry active histone modifications, including H3K4me1, H3K4me3, and H3K27ac. Yet, the principles governing the boundaries of such regulatory elements are still poorly characterized. Alu elements are good candidates for a boundary function, being highly abundant in gene-rich regions, while essentially excluded from regulatory elements. Here, we show that the interval from the TSS to the first upstream Alu accommodates all H3K4me3 and most H3K27ac marks, while excluding DNA methylation. Remarkably, the average length of this intervals greatly varies in-between tissues, being longer in stem-and shorter in immune-cells. The very shortest TSS-to-Alu intervals were observed at promoters active in T cells, particularly at immune genes, correlating with RNA polymerase II transcription through the first Alu and accumulation of H3K4me1 signal on this first Alu. Finally, DNA methylation at first-Alus was found to evolved with age, regressing from young to middle-aged, then recovering later in life. Thus, the first Alus upstream of TSSs appear as dynamic boundaries marking the transition from DNA methylation to active histone modifications at regulatory elements, while also participating in the recording of immune gene transcriptional events by positioning H3K4me1-modified nucleosomes.

## Introduction

In eukaryotes, transcription by RNA polymerase II (RNA Pol.II) is controlled by promoters, located just upstream of the transcribed region of genes, and by enhancers, eventually located at a distance. These regulatory elements (REs) are landing pads for transcription factors. They are also sites of bi-directional transcription initiation, enhancers producing short, unstable “eRNAs” in both directions, while transcription elongates in at least one direction at promoters (1). Finally, enhancers and promoters are sites of weakly positioned nucleosomes carrying specific histone modifications, including histone H3 lysine 27 acetylation (H3K27ac) enriched at enhancers, histone H3 lysine 4 tri-methylation (H3K4me3) enriched at promoters, and histone H3 lysine 4 mono-methylation (H3K4me1) enriched at enhancers and upstream of the H3K4me3 signal at promoters (2).

Large genome-wide projects such as Fantom or the NIH Roadmap Epigenomics Mapping have provided extensive information on location and tissue-specificity of REs (3, 4). In particular, these data have pinpointed how extensively the enhancer landscape evolves during development, with embryonic enhancers being frequently silenced during cell differentiation and replaced by other more tissue-specific enhancers at other locations (5).

This plasticity raises the question of the positioning of REs. Obviously, enhancer positioning relies largely on the presence of binding sites for transcription factors, that in turn recruit the RNA Pol.II, the histone modifying enzymes, and the chromatin remodeling complexes. Yet, transcription factor binding sites are poor indicators of RE boundaries, as their sequences are generally short and therefore also present outside of REs (6). The activity of such ectopic binding sites is controlled by DNA methylation at CpG dinucleotides, limiting transcription initiation to genuine promoters and enhancers (7). The repressive effect of DNA methylation is mediated by recruitment of methyl-binding proteins and associated histone deacetylases, maintaining a condensed local chromatin environment (8). In parallel, DNA methylation can also interfere with recruitment of histone acetyltransferases and, in some cases, with recruitment of transcription factors, particularly when a CpG is included in the recognized sequence (9, 10). However, DNA methylation does not entirely solve the issue of RE boundaries, as the targeting of DNA methylases remains poorly characterized, leaving unsolved the question on how is positioned the transition from unmethylated REs to the methylated surroundings (11, 12).

Another characteristic of REs is their depletion in transposable elements. Occasionally such elements will contribute with DNA binding sites, and they are considered as important drivers of evolution in the field of gene regulation (13–17). Yet, transposable elements are predominantly excluded from REs, and most sites of transcription initiation are located in repeat-free regions (18–20). In parallel, the DNA of transposable elements is subject to extensive methylation, preventing them from damaging the genome by novel insertions (21). There is therefore a likely link between absence of DNA methylation and absence transposable elements within REs.

In humans, Alu elements are the most successful of all mobile elements, present in more than 1 million copies and contributing almost 11% of the genome. They also contribute more than 25% of the CpG di-nucleotides (22). The sum of all Alu elements, previously referred to as the “Alu-ome”, is therefore an abundant matrix for DNA methylation, possibly regulated in a tissue-specific manner (23, 24). Counterintuitively, Alu elements are particularly abundant in euchromatin (or actively transcribed chromatin), where most genes are also located. This was discovered decades ago as Alu *in situ* hybridization labelling coincided with negative Giemsa staining (R-bands) on metaphase chromosomes (25). More recently, it was shown that the regions of positive Giemsa staining (G-bands) were matching Lamin A-associated Domains (or LADs), that allow heterochromatin to concentrate at the periphery of the nucleus by interacting with the nuclear lamina (26, 27). DNA sequencing has further confirmed the low Alu density within LADs as compared to the gene-rich transcriptionally-active interLADs (28). Consistent with a positive role of Alu elements in transcription, Alu elements were also reported to be enriched in H3K4me1, that, as mentioned above, is abundant at both enhancers and promoters (13). Finally, we note that Alu elements function as sites of nucleosome positioning, that may provide another avenue to transcriptional regulation (29, 30).

The exclusion of Alu elements from sites of transcription initiation, their propensity to be methylated, and their ability to position nucleosomes, prompted us to examine their potential role in setting the boundaries of REs. We find that the first Alu encountered by the RNA Pol.II during promoter and enhancer transcription is an inflection point for several epigenetic marks, delineating the decline of H3K4me3, while initiating the upstream DNA-methylation landscape. In a subset of tissues, particularly of hematopoietic lineage, we further observed a preferential positioning of REs in very Alu-dense regions, resulting in transcription start sites being very close to the first Alu. This TSS-to-first-Alu proximity further correlated with increased positioning of H3K4me1 signal at the first-Alu. Observation of data from newborn, middle-aged, and long-lived donors suggested that this H3K4me1 positioning participated in keeping a trace of earlier episodes of transcriptional activity at genes involved in immunity. Finally, observation of DNA methylation in the three age-groups provided evidence for a dynamic in the boundary function of Alu elements, possibly as a mechanism controlling access to upstream transcription factor binding sites.

## Material and Methods

### Data download

RNA-seq fastq files were downloaded from the Gene Expression Omnibus (GEO) and Sequence Read Archive (SRA) NCBI resources using the SRA toolkit (http://ncbi.github.io/sra-tools/). Accession numbers: GSE118106 (embryonic stem cells), GSE111167 (Jurkat T cells), GSE65515 (donor CD4+ T cells, RNA-seq and MeDIP data), GSE58638 (HCT116 H3K4me3 ChIP-seq), GSE152144 (HCT116 CXXC1 ChIP-seq), and GSE94971 (DamID LaminB1/Dam profile for resting Jurkat cells, per DpnI fragment, quantile normalized, smoothened in 60-fragment windows (∼15Kb) with a 1-fragment shift, in bigwig format). Chromatin states bed files (15-state model) as well as H3K4me3 and H3K27ac histone-modification bigwig files mapped on human genome version hg19 were fetched on the NIH Roadmap Epigenomics Mapping Consortium server (https://egg2.wustl.edu/roadmap/web_portal/imputed.html). Bed file with CAGE peaks for human samples (phase1 and 2 combined) were fetched on the Riken FANTOM5 server (https://fantom.gsc.riken.jp/5/data/). Mapping of Alu elements on the hg19 version of the human genome were extracted from RepeatMasker (https://www.repeatmasker.org/).

### RNA-seq Mapping

Mapping was carried out with STAR (v2.6.0b) (parameters: --outFilterMismatchNmax 1 -- outSAMmultNmax 1 --outMultimapperOrder Random --outFilterMultimapNmax 30) (31). The reference genomes were hg19 homo sapiens primary assembly from Ensembl. The SAM files were converted to BAM files and sorted by coordinate using samtools (v1.7) (32).

### MeDIP-seq Mapping

To maximize MeDIP read mapping at repetitive elements we took advantage of the data being from paired-end sequencing, aligning each mate separately and we generated a pipeline requiring only unambiguously of one of the two mates. Specifically, reads were mapped to the human genome (hg19 homo sapiens primary assembly from Ensembl) using bowtie2 (v2.3.4) (33) (parameters: -N 0 -k 1 –very-sensitive-local). The SAM files were then converted to BAM files and sorted by coordinate using samtools (v1.7) (32). We then selected reads with a MAPQ equal or higher than 30 and then re-associated with their mate (that may have a low MAPQ score).

### BigWig files, heatmaps, profiles, and data quantification

Bigwigs files were generated from .bam files with bamCoverage (parameter: --normalizeUsing CPM) from Deeptools (v3.1.3) (34). For H3K4me1, the .bam files were obtained by converting tagAlign ChIP-seq files from the Roadmap Epigenomics project repository using the bedToBam function from samtools (v1.7) [2].For Alu element distribution, .bam files were obtained by extracting in .bed file format, entries annotated “Alu” in the “RepFamily” field from RepeatMasker, then converting the .bed to .bam files with bedtobam from the bedtools package (v2.27.1) from the Quinlan laboratory (http://quinlanlab.org). Heatmaps and profiles were generated with Deeptools (v3.1.3). Matrices were generated with computeMatrix followed by plotProfile or plotHeatmap as appropriate. When indicated in the figure captions, matrices were built on a narrow region centered on the first-Alu (parameter: -b 500 -a 50, relative to the 3’ end of the first-Alu), then heatmaps were plotted on a wider region using the parameter “--sortRegions keep”. Clustering was performed using the Kmean algorithm, either on narrow or wider regions as indicated in the captions. Read quantification was carried out with featureCounts (v1.6.1) from the Subread suite (35).

### Data visualization

The Integrative Genomics Viewer software (IGV) was used to examine specific loci (36). The R/Bioconductor package karyoploteR (37) was used to plot whole genomes with <1kbCIAs and IRIS immune genes (38).

### Similarity index

To compare distribution of Alu elements to that of promoters and enhancers: (1) entries annotated “Alu” in the “RepFamily” field from RepeatMasker were extracted in .bed. (2). Bed files with either promoter or enhancer regions were generated by extracting regions annotated with the mnemonics “1_TssA” or “7_Enh”, respectively. Jaccard indexes were calculated with bedtools 2.27.1.

### Gene ontology

Identification of genes in the neighborhood of CIAs of indicated sizes was carried out with GREAT (39). KEGG pathway analysis on genes with peaks of H3K4me1 on their promoter first-Alu in all tissues under scrutiny was carried out with Enrichr (40). Bar-graphs show -log10 of the binomial p value.

## Results

### Tissue-specific distribution of regulatory elements in “the Alu-ome”

To explore the possible function of Alus at the boundaries of RE, we first re-investigated how promoters and enhancers locate relative to these retroelements. To that end, we mined chromatin state data from the NIH Roadmap Epigenomics Mapping Consortium. These “chromatin states” were predicted by combining data on histone modifications, DNA methylation, and chromatin accessibility in 127 different human tissues (4). From these data, we extracted regions annotated either as transcription start sites (TssA) or enhancers (Enh) in all the tissues. To compared the distribution of these REs to that of Alu elements, we used the Jaccard index. This index, also known as the similarity index, is defined as the size of the intersection divided by the size of the union of the sample sets. As a control, we also calculated the Jaccard index between the two types of REs and randomly selected Alu-free regions (average of thousand iterations). The score shown for each tissue is the Jaccard index (REs vs. Alus) divided by the control Jaccard index (REs vs. random).

For promoters, the relative Jacquard index systematically remained below 1 in all tissues, establishing a clear counterselection of Alu elements inside these REs (Sup. Figure 1A). In contrast, for enhancers, the score varied extensively from one tissue to the other, Alu elements being positively selected at enhancers from some tissues, while counter-selected in others (Figure 1A). The quartile with the strongest counterselection included mostly stem, fetal, and brain tissues. In contrast, the quartile with strongest positive selection exclusively contained hematopoietic tissues or tissues rich in hematopoietic cells (placenta). Visual examination of the distribution of regions annotated as enhancers with a genome browser confirmed their more frequent overlap with Alu sequences in high-scoring tissues as compared to tissues with counterselection (see example Figure 1B).

**Figure 1:**
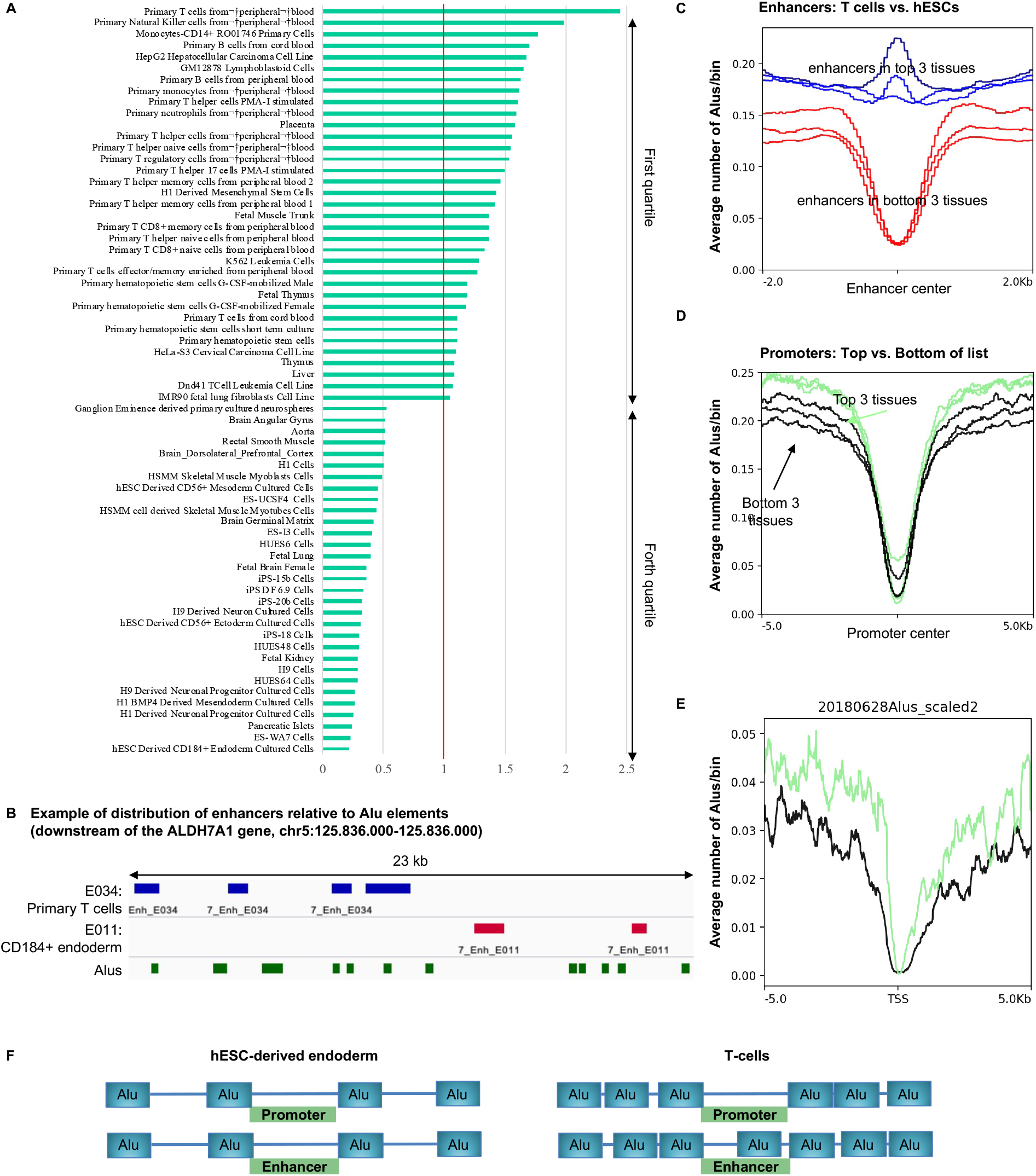
Tissue-specificity in the positioning of Alu elements relative to REs. (A) Comparison of regions annotated “Alu” in RepeatMasker with the regions annotated “7_Enh” in the 15 core marks model of the Epigenomic Roadmap consortium. For each tissue, the Jaccard index comparing enhancers to Alus is divided by the average Jaccard index (1000 iterations) obtained when comparing enhancers to randomly selected genomic locations (of the same sizes as the Alus). Only the tissues in the first and the last quartiles with the highest and the lowest scores are shown. (B) Screen capture from IGV representing an example of distribution of enhancers relative to Alu elements in the tissue with the highest and the lowest score, respectively E034 T cells and E011 hESC-derived endoderm. (C-D) Plots representing the average distribution of regions annotated “Alu” in RepeatMasker relative to enhancers or promoters from the indicated tissues. (E) Transcriptome data from either T cells (green) or human embryonic stem cell (black) were used to identify marker genes for each of the two cell types. Plots represent the average distribution of regions annotated “Alu” in RepeatMasker relative to the transcription start site (TSS) of the two sets of marker genes. (F) Schematic of the deduced distribution of Alu elements relative to enhancers and promoters in E011 hESC-derived endoderm and in E034 T cells.

In order to also apprehend density in Alus at regions framing REs, we further plotted the average distribution of Alu elements relative to enhancers and promoters for the three top-and bottom-scoring tissues (Top: E034 primary T cells from peripheral blood, E046 primary natural killers from peripheral blood, and E124 monocytes-CD14+ RO01746 primary cells; Bottom: E011 hESC-derived CD184+ endoderm cultured cells, E002 ES-WA7 cells, and E087 pancreatic islets). These graphics first confirmed that the top tissues accommodated Alu elements within the boundaries of their enhancers, while, in the bottom tissues, enhancers appeared as Alu-depleted valleys (Figure 1C). Among the top tissues, E034 was particularly noticeable as the only tissue with a clear peak of Alu density associated with the average enhancer (Figure 1C, dark blue profile). Examining the surroundings of the enhancers further revealed that top-tissue enhancers were in more Alu-dense environments than were bottom-tissue enhancers (Figure 1C, blue profiles always above red profiles). Similarly, at promoters, that were Alu valleys in all tissues, the flanking regions of the top-tissues showed a higher density in Alu elements than did the bottom-tissues (Figure 1D, green profiles always above black profiles).

To confirm this difference with independent data, we used transcriptomes from either T cells (41) or human embryonic stem cell (42) to identify marker genes for each of the two cell types. Plotting of the average Alu density over these two series of genes confirmed that promoters of T cell specific genes were in average located in more Alu-dense chromosome regions than stem-cell-specific genes (Figure 1E).

Together, these observations documented that regions annotated as promoters by the Epigenomic Roadmap Consortium strictly evade Alu elements, while these elements were eventually accommodated inside regions annotated as enhancers, particularly in hematopoietic tissues. In parallel, the analysis suggested that Alu-density at the boundaries of active RE also shows tissue-specific variations, being high at REs active in T cells, and lower at those active in several lines of stem cells (schematic Figure 1F).

### Immune cell regulatory elements locate to regions of high Alu-density

We next examined more systematically the Alu density in the neighborhood of REs active in each tissue from the NIH Roadmap Epigenomics Mapping Consortium. As a reporter of Alu density, we used the interval in-between two Alu elements that, in average, will shorten as a function of the increased density (Figure Sup. 2A). As REs may eventually include Alu elements and therefore be overlapping with more than one Alu-to-Alu interval, we identified for each RE, the Alu-to-Alu interval containing the site of transcription initiation (see schematic Figure 2A). For this, we relied on the Fantom5 repository of transcription start sites (TSSs) consolidated from 975 libraries of Cap Analysis Gene Expression (CAGE) (3). This allowed us to identify Alu-to-Alu intervals containing at least one CAGE peak, henceforth referred to as “CAGE-containing InterAlus” (CIAs). We then crossed these CIAs with enhancers and promoters from the NIH Roadmap Epigenomics Mapping Consortium data to identify “transcriptionally active” CIAs in each of the 127 tissues (see explanatory schematic Sup. Figure 2B). When the tissues were ranked as a function of the median size of their transcriptionally active CIAs, the order was remarkably similar to that observed in Figure 1A, and T cells clustered among the tissues with the lowest median size CIAs, while embryonic cells displayed the highest ones (Figure 2B). This was illustrated graphically by plotting the average Alu distribution over active CIAs in the 3 top-and bottom-scoring tissues (Figure 2C, blue profiles always above red profiles). A bar-graph further visualized the CIA size distribution in E034 primary T cells from peripheral blood (shortest CIAs) and in E011 hESC-derived CD184+ endoderm cultured cells (longest CIAs – Sup. Figure 2C). The difference in average CIA size was particularly exacerbated when focusing on enhancers in top-and bottom-tissue (E034 and E011 - Figure 2D).

**Figure 2:**
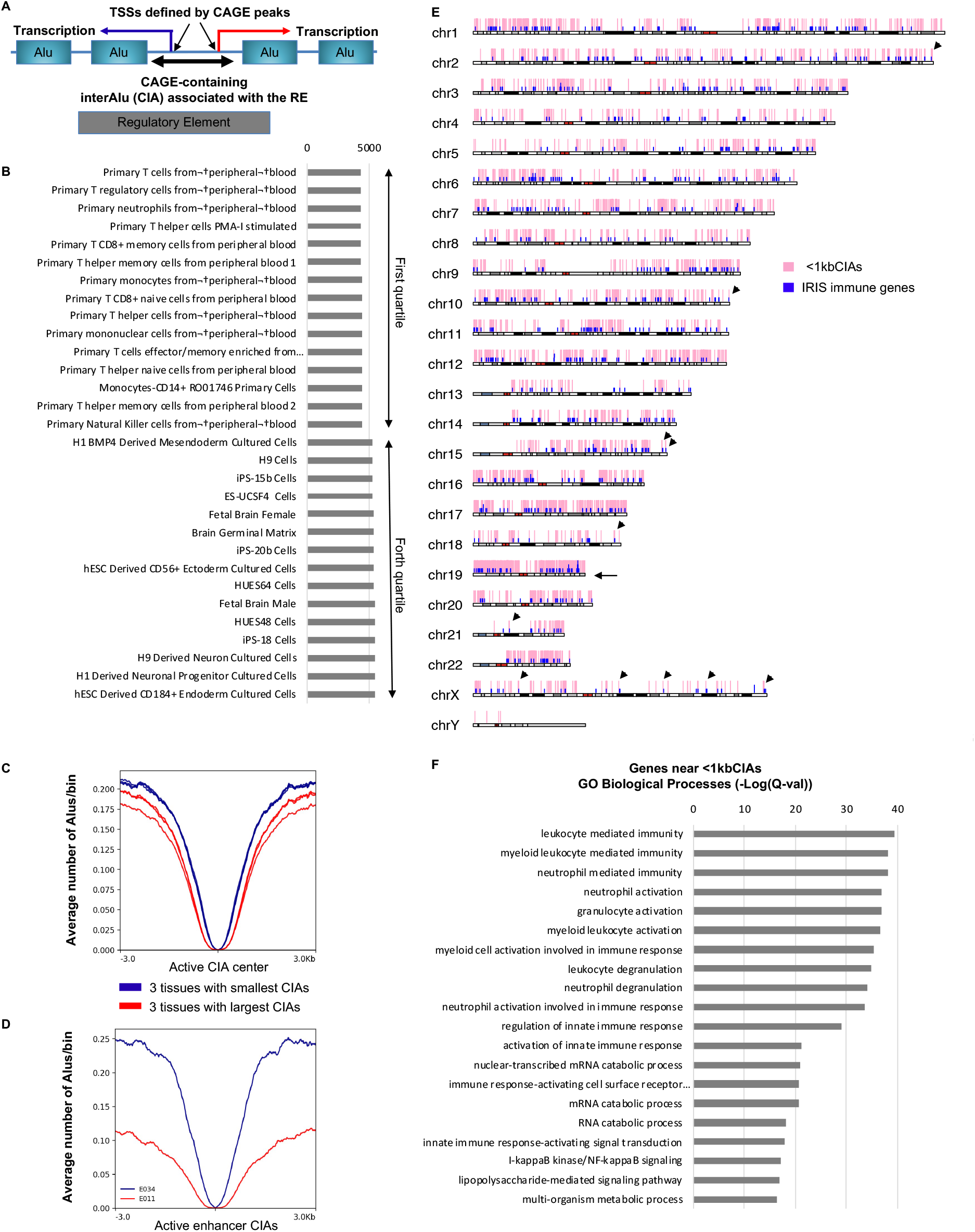
RE in regions of high Alu density are a specificity of immune genes. (A) Schematic defining a CAGE-containing interAlu (CIA) as a genomic region framed by two Alu elements and hosting at least one Fantom5 CAGE peak. If the site of transcription initiation is active in a given tissue, a CIA is expected to overlap with a regulatory element a defined by the NIH Roadmap Epigenomics Mapping Consortium (grey box). (B) For each of the 127 tissues annotated by the Epigenomic Roadmap consortium, the average size of CIAs intersecting with regions designated as either “1_TssA” or “7_Enh” was calculated. The bar graph shows the 15 tissues with lowest and highest average CIA size. (C) Plot representing the average distribution of regions annotated “Alu” in RepeatMasker relative to CIAs for the 3 tissues with either the shortest (blue profiles) or the longest (red profiles) average CIA sizes. (D) Plot representing the average distribution of regions annotated “Alu” in RepeatMasker relative to enhancers overlapping with CIAs in either E034 T cells (shortest average CIA size – blue profile) or E011 hESC-derived endoderm (longest average CIA size – red profile). (E) Karyoblot representing the position of immune genes as defines by the IRIS collection and of CIAs less than 1kb in length (<1kbCIAs) in the indicated colors. (F) Genes located in the neighborhood of <1kbCIAs were identified using GREAT. The list of genes was then analyzed for enrichment in GO terms. Histogram shows the false discovery rate as -Log(Q value) as calculated by GREAT.

Next, to examine the link between Alu density and gene expression independently of the NIH Roadmap Epigenomics Mapping Consortium data and its associated chromHMM chromatin-state prediction algorithm, we monitored the distribution of CIAs having a length of less than 1kb (4981 intervals in total, referred to as <1kbCIAs). In total, we identified 64,896 CIAs, with a median size of 4.2kb. Thus, the <1kbCIAs represented approximately the 10% shortest CIAs. The density in <1kbCIAs was particularly high on chromosome 19 (Sup. Figure 2D), consistent with this chromosome being particularly rich in Alu elements (43). Interestingly, this chromosome is also enriched in genes involved in immunity (38). This prompted us to plot <1kbCIAs together with immune genes as defined by the IRIS collection (44). This graphic confirmed the high density on chromosome 19, while also revealing that <1kbCIAs frequently coincided with immune genes at many sites along the chromosome (Figure 2E, arrows indicate examples).

To quantify this apparent colocalization, we identified genes neighboring the <1kbCIAs and analyzed the result for GO term enrichment (using GREAT at default setting - Proximal: 5kb upstream, 1kb downstream, plus Distal: up to 1000kb) (39). This approach identified 68 significantly enriched pathways, including 36 related to immunity or immune cell, the top scoring pathways including leukocyte mediated immunity, myeloid leukocyte mediated immunity, and neutrophil mediated immunity with FDR Q-values in the range of 10^−38^ (Top20 in Figure 2F). Pathways related to RNA metabolism were also abundantly represented (8 pathways). When, as a control, the ensuing 4981 larger CIAs (ranging from 1000 to 1525 nucleotides in size) were analyzed in the same way, neighboring genes were associated with mostly unrelated pathways (14 pathways) with best FDR Q-values in the range of 10^−9^ (Sup. Figure 2E). Likewise, genes in proximity of a series of 4981 intervals centered on the median CIA length (ranging from 3908 to 4615 nucleotides) also identified unrelated pathways (11 pathways with best FDR Q-values in the range of 10^−5^ - Sup. Figure 2F).

Together, these two independent approaches to probed the usage of REs located in Alu-dense regions respectively identified either cell types from the hematopoietic lineage, or genes involved in immunity, strongly suggesting that Alu-dense neighborhoods provide strategic advantages to organismal defense mechanisms. The possible benefits for gene regulation will be explored below.

### T cells frequently position H3K4me1 histone marks at the Alu preceding a TSS

To gain insight in the possible function of short CIAs, we examined the distribution of the H3K4me1 histone mark over Alu elements adjacent to TSSs. This histone modification is enriched at enhancers, but also at promoters where it locates immediately upstream of the H3K4me3 signal (45). H3K4me1 is also enriched at Alu elements located in the proximal upstream region of genes in T cells (13). Finally, this histone mark was associated with a memory function, accumulating at promoters as a consequence of temporary transcriptional activity (46).

First, we explored the distribution of H3K4me1 at promoters, anchoring the profiles on the 3’ end of the first Alu after the TSS in the orientation of “upstream antisense RNA” (uaRNA)-transcription *i*.*e*. opposite to gene-transcription (henceforth referred to as the “first-Alu” - see schematic Figure 3A). In E034 T cells, these profiles revealed the expected H3K4me1 peak on the first-Alu (Figure 3B). This peak was not observed when using H3K4me1 data from E011 CD184+ endoderm (Figure 3C). At note, in both profiles, there was a loss off signal over the first-Alu, a predictable consequence of the filtering of multimapping reads over repeated sequences during the alignment procedure.

**Figure 3:**
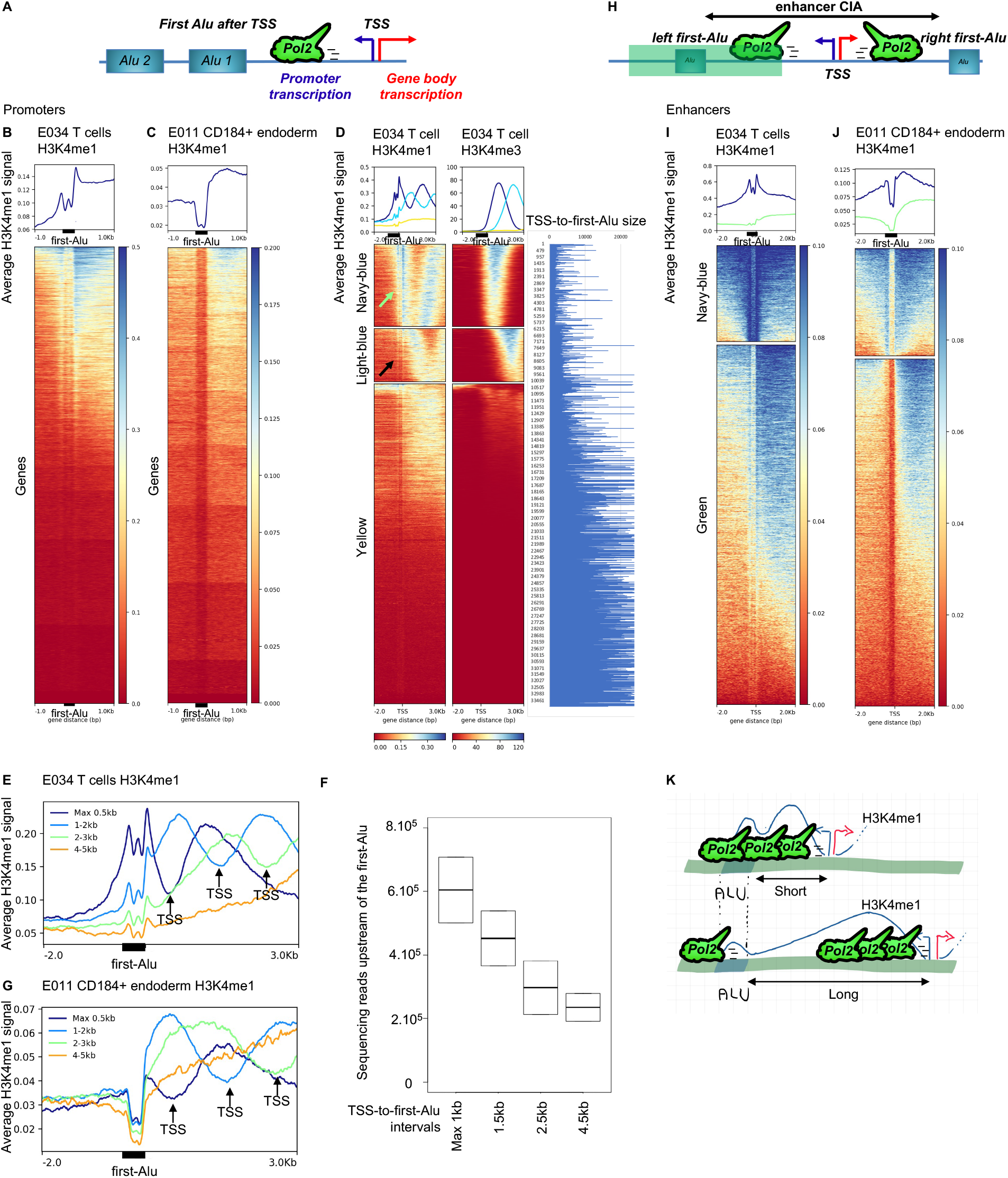
Peaks of H3K4me1 at first-Alus is favoured by proximity to the TSS. **Promoters:** (A) Definition of the first Alu element encountered by the RNA POLII after initiation at the TSS in the orientation of promoter transcription. (B-C) Heatmaps of H3K4me1 signal from E034 T cells and E011 hESC-derived endoderm as indicated at first-Alus from promoters. The heatmaps are anchored on the 3’ end of the first-Alu. The position of the 5’ end of the black box symbolizing the first-Alu is an approximation. The first-Alus are sorted in the order of decreasing signal from -500 nts to +50 nts relative to the 3’ end of the first-Alu. (D) Promoter first-Alus where clustered in 3 clusters using the k-mean algorithm, based on the H3K4me3 signal on a region spanning from -2000 to +3000 nts relative the 3’ end of the first-Alu. Using these clusters (each associated with a color), heatmaps with H3K4me1 and H3K4me3 signals were plotted in parallel, and sorted in the order of decreasing H3K4me3 signal. The rightmost bar graph shows for each first-Alu, the sizes in nucleotides of the TSS-to-first-Alu intervals. (E, G) For the indicated tissues, profiles of H3K4me1 signal anchored on the first-Alus, for TSS-to-first-Alu intervals having a length of either less than 0.5kb, 1 to 2kb, 2 to 3kb, or 4 to 5kb. The indicated positions of the TSS in each size-range is deduced from the inflexion point in the H3K4me1 signal. (F) Using RNA-seq data from Jurkat T cells (N=3 for each condition), reads mapping withing 500 nts upstream of the first-Alu 5’-end were quantified for TSS-to-first-Alu intervals having the indicated length. **Enhancers:** (H) Definition of left and right first-Alus (in the orientation of the genome) at enhancers. The region covered by the heatmaps (I) and (J) is shaded in green. (I-J) Distribution of the H3K4me1 signal over the “left first Alus” from CIAs overlapping with enhancers in the indicated tissues. The heatmaps are anchored on the 3’ end of the Alu elements and clustered and sorted based on the signal present in the interval -500 nts to +50 nts relative to the 3’ end of the Alus. The position of the 5’ end of the black box symbolizing the Alu is an approximation. (K) Interpretation: (Top diagram) when the first-Alu is close to the TSS, it is run through by the RNA PolII, favouring positioning of the H3K4me1 signal. (Bottom diagram), when the first-Alu is distant from the TSS, it is reached only occasionally by the RNA PolII and the H3K4me1 signal.

To investigate a possible role for transcription in the positioning of H3K4me1 signal over the first-Alu, we next used E034 T cell H3K4me3 signal as a surrogate reporter of transcriptional activity. K-means clustering allowed to segregate a set of promoters at which the H3K4me3 signal was edging at the first-Alus (Figure 3D, navy-blue clusters). This cluster, also enriched in the shortest TSS-to-first-Alu intervals (Figure 3D, leftmost bar graph), displayed a clearer positioning of the H3K4me1 signal than did the subsequent cluster encompassing promoters with H3K4me3 signal located away from the first-Alu (Figure 3D, compare black and green arrows). This suggested that promoters with short TSS-to-first-Alu intervals were prone to position H3K4me1 at their first-Alu when transcriptionally active.

To further investigate this possibility, we segregated promoters in bins as a function of the size of their TSS-to-first-Alu interval, and then plotted the average H3K4me1 distribution profile, anchored on the 3’ end of the first-Alus, for each size bin. This approach revealed a clearly defined H3K4me1 peak at the shortest TSS-to-first-Alu intervals, gradually decreasing in intensity as intervals were lengthening (Figure 3E). This peak segregated best from the main signal when the TSS-to-first-Alu intervals were in the size ranges 1-to-2Kb and 2-to-3Kb (Figure 3E, light-blue and green profiles). Importantly, increased TSS-to-first-Alu size also correlated with a decreased transcription of the first-Alu, as documented by quantification of sequencing reads mapping to the region immediately after the first-Alu in the orientation of promoter transcription (Figure 3F). This suggested a correlation between transcription through the first-Alu and positioning of the H3K4me1 signal at the first-Alu.

First-Alu H3K4me1 peaks were not observed when using H3K4me1 data from E011 hESC-derived CD184+ endoderm (Figure 3G). This prompted us to systematically examine the 1-to-2Kb and 2-to-3Kb TSS-to-first-Alu intervals in the 127 tissues from the NIH Roadmap Epigenomics Mapping Consortium, to identify tissues positioning H3K4me1 peaks at their first-Alus. While the outcome was sometimes ambiguous, we identified clear H3K4me1 positioning in several non-T cell data sets, including several muscle tissues (Figure S3A).

Finally, to determine whether enhancers also generated peaks of H3K4me1 signal at Alu elements close to transcription start sites, we selected CIAs overlapping with E034 enhancers, then examined the signal profile over the Alus at the ends of these CIAs (left first-Alu in the orientation of the genome, Figure 3H). Clustering based on the intensity of the H3K4me1 signal over these Alus allowed identifying a series of enhancers with peaks localized at that position (Figure 3I). Unlike what we observed at promoters, these events appeared internal to enhancers, with abundant H3K4me1 signal on both sides of the Alu. The complementary cluster of enhancers, with lower levels of H3K4me1, had a more promoter-like distribution, with the signal mostly on the inside of the CIA. The same analysis on E011 enhancers showed that in that tissue, the CIA boundaries were not inflection points for the H3K4me1 signal (Figure 3J).

Together, these observations confirmed that H3K4me1 signal is frequently positioned at promoter-proximal Alu elements in T cells. Furthermore, they show that this positioning is favored by the high Alu density (or short TSS-to-first-Alu intervals) characteristic of the environment of REs active in T cells, a characteristic that promotes serendipitous transcription of the Alu elements by the RNA PolII (Schematic Figure 3K). Finally, exploring other tissues revealed that the positioning of the H3K4me1 signal at first-Alus is not restricted to T cells, and may also be promoted by cues other than TSS-to-first-Alu interval length, for example, high levels of promoter activity.

### The first-Alu element is an inflection point for DNA methylation and H3K4me3 marks

In addition to their impact on the H3K4me1 signal, Alu elements have been largely studied for their high level of DNA methylation, an epigenetic modification associated with transcriptional repression. To gain insight in the possible consequences of Alu density on DNA methylation at T cell REs, we mined a MeDIP data set from donors at three different ages, either newborn, middle-aged adults, or long-lived (41). Graphic examination of the MeDip data showed frequent peaks of DNA methylation over the more recent AluY and AluS members of the Alu family (Sup. Figure 4A). Peaks over the older AluJ members, having lost most of their CpG content, were rarer. This was confirmed genome-wide by heat maps reporting DNA methylation at 50,000 randomly selected Alu elements from each family (Sup. Figure 4B). From these observations, we concluded that this data set agreed with expected Alu methylation patterns.

**Figure 4:**
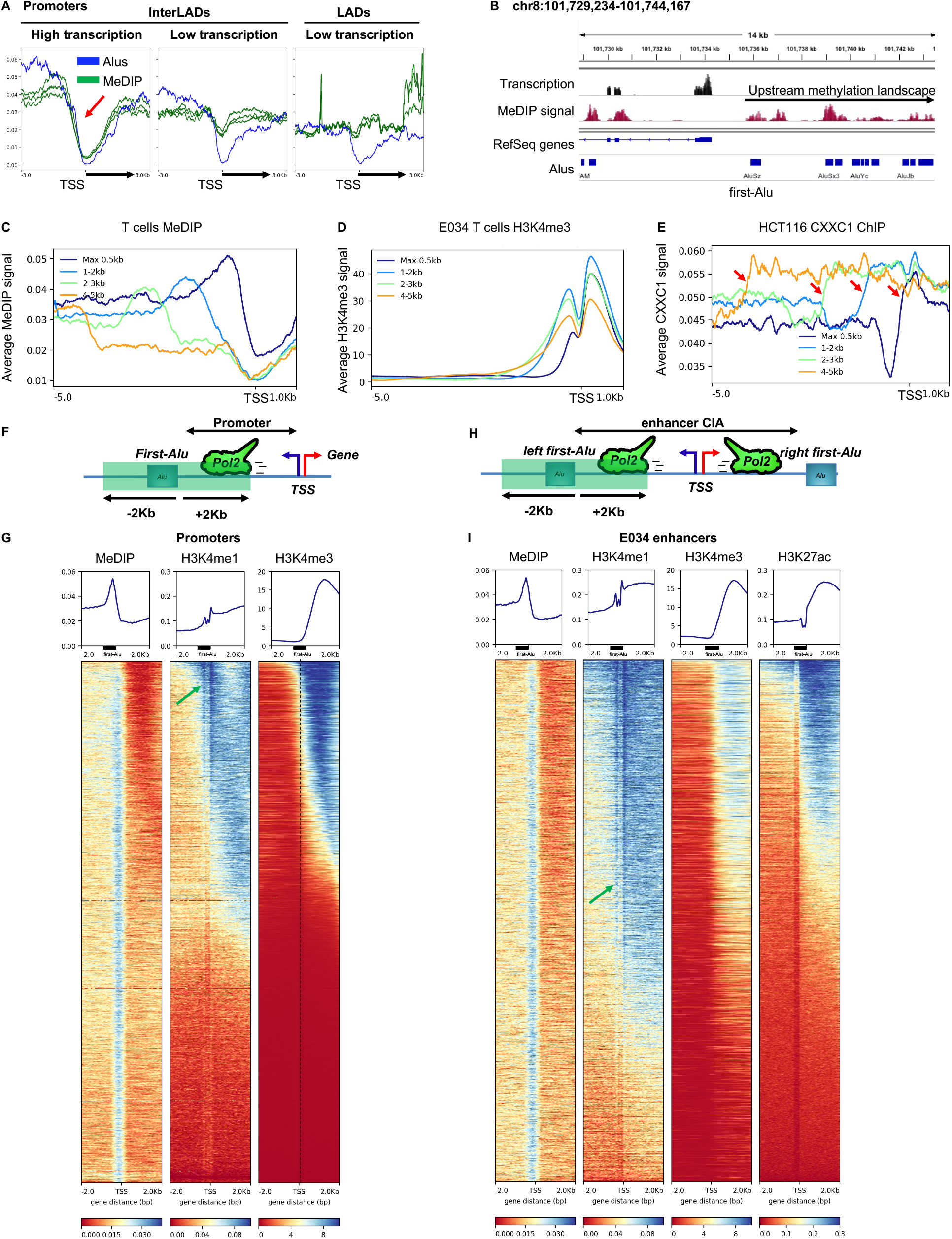
First-Alus are inflection point for epigenetic marks at regulatory elements. (A) Profiles of Alu element distribution and MeDIP signal (N=3) at 1000 genes with either the highest or the lowest expression levels (based on RNA-seq data from the middle-aged donors – N=3) and located in interLADs, and at 1000 genes locating to LADs as indicated. Profiles are anchored on the transcription start site (TSS) of the genes. Black arrows indicate the transcribed regions of genes. Red arrow indicate coupling of Alu distribution with DNA methylation. (B) Screenshot from IGV – Example of MeDIP signal distribution at a transcriptionally active gene. (C-E) Average distribution profiles anchored on the TSS, for either MeDIP, H3K4me3, or CXXC1 as indicated. Profiles are for TSS-to-first-Alu intervals having a length of either less than 0.5kb, 1 to 2kb, 2 to 3kb, or 4 to 5kb. (F) Schematic depicting the region explored in ensuing promoter heatmaps, anchored on the 3’ end of the first-Alu and extending 2Kb in each direction (region shaded in green). (G) Promoters: heatmaps with MeDIP, H3K4me1, and H3K4me3 signals were plotted in parallel and sorted in the order of decreasing H3K4me3 signal. The green arrow shows dense H3K4me1 signal. (H) Schematic depicting the region explored in ensuing enhancer heatmaps, anchored on the 3’ end of the left first-Alu (in the orientation of the genome) and extending 2Kb in each direction (region shaded in green). (I) Enhancers: heatmaps with MeDIP, H3K4me1, H3K4me3, and H3K27ac signals were plotted in parallel and sorted in the order of decreasing H3K4me3 signal. The green arrow shows dense H3K4me1 signal.

We then focused on the middle-aged donors, considered as the most representative of adulthood, and with this dataset, we examined the impact of Alu density on DNA methylation at promoters. To account for the link between methylation and transcription, we examined separately the 1000 most and least expressed genes. Genes located in LADs were also put in a separate list, as LADs are overall regions of low Alu density (see example at the short arm of chromosome 2, Sup. Figure 4C). Plotting the average MeDIP signal at the transcriptionally most active genes yielded a profile remarkably similar to that of the Alu distribution (compare blue and green profiles, left panel, Figure 4A). In contrast, at genes displaying low expression in T cells or at genes located in LADs, the MeDIP signal was uncoupled from the Alu content, with average levels essentially unaffected by the Alu-valley surrounding the TSS (central and right panels, Figure 4A). These profiles strongly suggested that, in the absence of negative regulation of transcription, density in Alu elements was a major determinant of DNA methylation at regions located upstream and downstream of active promoters. Reciprocally, these observations also suggested that methylation intended for negative regulation of genes depended on mechanisms independent of Alu elements.

Examination of several active promoters with a genome browser suggested that the methylation landscape upstream of transcribed genes was initiated at the first-Alu (see example of the PABPC1, Figure 4B). To examine more systematically whether the position of first-Alus influenced that of epigenetic marks at promoters, we segregated promoters in bins as a function of the size of their TSS-to-first-Alu interval as described in Figure 3. Then, we plotted the average distribution of T cell MeDIP, H3K4me1, and H3K4me3 signals anchored on the TSS (Figures 4C, 4D, and Sup. 4D). These profiles showed that increasing TSS-to-first-Alu distance directly impacted on the distribution of the epigenetic marks, each drifting towards more upstream promoter regions as TSS-to-first-Alu distances were increasing. A similar drift was also observed when plotting E011 CD184+ endoderm H3K4me1 and H3K4me3 signals, suggesting that an effect of first-Alus on the positioning of epigenetic marks was not a tissue-specific phenomenon (Figures Sup. 4E and Sup. 4F).

To gain mechanistic insight on the apparent first-Alu boundary effect, we examined the distribution of CXXC1, a subunit of the COMPASS methyltransferase, responsible for most H3K4 tri-methylation (47). This zinc-finger protein binds unmethylated CpGs and thereby participate in the targeting of the H3K4me3 mark to active promoters. As high quality CXXC1 ChIP data was not available in T cells, we examine the distribution in HCT116 colon carcinoma cells after verifying that distribution of the H3K4me3 signal relative to first-Alus was similar to that observed in T cells (Sup. Figure 4G). In the HCT116 cells, the point at which CXXC1 recruitment returned to background levels moved upstream as the TSS-to-first-Alu interval increased in length (Figure 4E). This was consistent with first-Alu DNA methylation being a driver of the boundary effect, by interfering with COMPASS recruitment.

To further visualize the boundary effect of the first-Alus, we plotted parallel heatmaps with MeDIP, H3K4me1, and H3K4me3 signals centered on the 3’ end of the first-Alus (see schematic Figure 4F). These heatmaps, sorted in the order of decreasing H3K4me3 signal, confirmed the asymmetry of the MeDIP signal intensity relative to the first-Alu at transcriptionally active promoters (left panel, Figure 4G, using H3K4me3 as a reporter of transcriptional activity). The heatmaps also confirmed a strong inflection in the H3K4me3 signal at the first-Alus, although H3K4me3 was reaching that point at only a subset of promoters (Figure 4G, right panel). Finally, H3K4me1, when reaching the first-Alu, displayed a peak of signal at that position, consistent with the observations described in Figure 3. But then, eventually, the H3K4me1 reached beyond the first-Alu, suggesting that the first-Alu was not a boundary for this modification (Figure 4G, arrow).

To also examine positioning of epigenetic marks at enhancer, we plotted heatmaps centered on the Alu elements located at the end of CIAs matching E034 enhancers (Schematic Figure 4H). In this series, the regions were sorted as a function of ascending H3K27ac signal (Figure 4I). As for the promoters, we observed a clearly asymmetric distribution of the MeDIP and H3K4me3 signal relative to the first-Alus. The H3K27ac signal was enriched but not strictly contained on the inside of the CIA. Finally, the H3K4me1 signal abundantly crossed the first-Alu boundary, a phenomenon that may explain the inclusion of Alu elements inside enhancers described in Figure 1. As observed at promoters, H3K4me1 crossing the first-Alu correlated with a peak of signal at that position (Figure 4I, arrow). Together, these observations indicated that at transcriptionally active RE, the first-Alu is a site of transition from DNA methylation to H3K4 trimethylation. As H3K4me3 is promoter-enriched, the mutual exclusion between this histone mark and first-Alus may explain why no overlap was observed between annotated promoters and Alu elements in Figure 1. Inversely, the spreading of H3K27 acetylation into first-Alus and the homing of H3K4 mono-methylation at these positions are compatible with an eventual overlap between Alu elements and annotated enhancers.

### Dynamics of epigenetic modifications at first-Alus throughout lifetime

The observations described above indicated that first-Alu methylation functioned as a barrier, interfering with upstream spreading of the H3K4me3 signal. To explore whether this barrier was dynamic, we took advantage of the high quality of the MeDIP data from the newborn, middle-aged, and long-lived donors used in Figure 4 (41). In LADs, overall displaying moderate transcription activity and low Alu density, we observed only small variations in the overall methylation of Alu elements between age groups, at the limit of significativity (Sup. Figure 5A). In contrast, in transcriptionally active interLADs, DNA methylation at these repeated elements decreased from infants to middle-aged, then increased from middle-aged to long-lived, in a context where overall methylation was either stable or moderately increased, as indicated by quantification of the MeDIP signal at randomly selected regions (Figure 5A). These variations were particularly exacerbated when examining the more recently integrated Alu family members, AluY and AluS (Figure 5B). Further examination of the differentially methylated regions (DMRs) identified in the initial study, confirmed that a large fraction of the sites having lost methylation at the transition from infant to middle-aged were enriched in Alu elements (Figure 5C, yellow arrow), while, at the transition from middle-aged to long-lived, Alu-enrichment was to be found in DMRs gaining methylation (Figure 5C, green arrow). These observations defined Alu elements as abundantly contributing to sites of differential methylation in T cells, loosing methylation during times of immune system maturation, then regaining methylation in late life.

**Figure 5:**
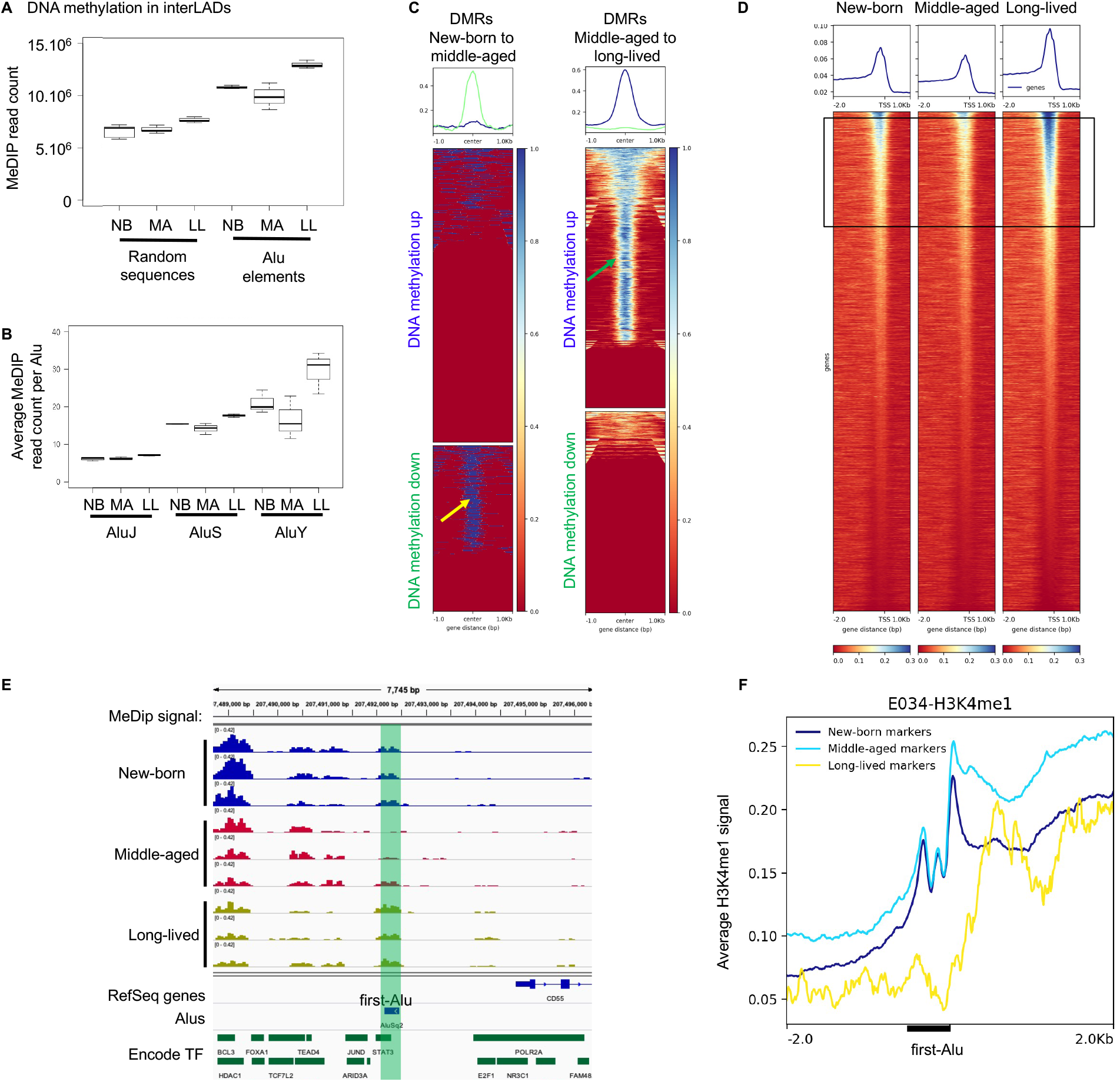
First-Alu DNA methylation and H3K4me1 peaking fluctuates with age. (A) Methylation inside Alu elements or randomly selected regions in interLADs at three different ages: counting of MeDIP reads either mapping to Alu elements located outside LADs or mapping to an identical set of non-Alu non-LAD regions in the indicated age-groups. NB: newborn, MA: middle-aged, LL: long-lived. (B) Counting of MeDIP reads mapping at either AluJ, AluS, or AluY family members in the indicated age-groups. (C) Heatmaps representing the Alu distribution relative to DMRs respectively undergoing increased or decreased DNA methylation when comparing new-borns to middle-aged or middle-aged to long-lived as indicated. (E) Screenshot from IGV – MeDip signal at the first-Alu upstream of the CD55 gene in each of the replicates at the indicated ages. Bottom track reports ENCODE sites of transcription factor binding. (F) Distribution profile of the H3K4me1 signal from adult T cells was plotted over the first-Alu at markers genes for new-born (2636 genes), middle-aged (2727 genes), long-lived donors (55 genes). (G-H) Model part 1: At promoters, the first-Alu sets the limit between the upstream DNA methylation landscape and the promoter-borne H3K4me3 signal. Unlike the H3K4me3 signal, the H3K4me1 crosses the first-Alu; this mark also accumulates at the first-Alu. At enhancers, DNA methylation and H3K4me1 distribute as described for promoters. H3K27 acetylation accumulates mostly in the interval between TSS and first Alu, but eventually spreads beyond the Alu boundary. (I) Model part 2: nucleosomes are poorly positioned on the promoter region, but well-positioned at the first-Alu (line 1). At promoters with long TSS-to-first-Alu intervals, the RNA PolII rarely reaches the nucleosomes positioned on the first-Alu and both H3K4me3 and H3K4me1 locates to poorly positioned nucleosomes. In contrast, when TSS-to-first-Alu intervals are short, the RNA PolII reaches the nucleosomes positioned on the first-Alu and favors their H3K4me1 modification. DNA methylation, by interfering with recruitment of the histone methyltransferase, then prevents transition from H3K4me1 to H3K4me3 at these nucleosomes and thereby spreading of the H3K4me3 mark (line 2). After the phase of activity, nucleosome replacement gradually erases most H3K4me1 marks. But on the first-Alu, where nucleosomes are stably positioned, the modification persist long after transcription has ceased. Due to the presence of these histone marks, previously active promoters with short TSS-to-first-Alu intervals are poised for later reactivation (line 3). (J) Model part3: when first-Alu methylation is low (*i*.*e*. affecting only a small fraction of the cell population), transcription factor binding sites upstream of the first-Alu are located in an area carrying H3K4me3 marks and allowing for transcription factor binding and/or activity. When first-Alu methylation is high, the active promoter region stops at the first-Alu, and upstream transcription factor binding sites are out of commission.

To test the impact of these variations on the boundaries of DNA methylation at promoters, we examined DMRs overlapping with first-Alus. Intersection of first-Alus with DMRs gaining methylation from middle-aged to long-lived was 2-fold more frequent than expected by chance (out of 9682 DMRs, 714 overlapped with first-Alus while only 361 overlapped with an identical number of randomly selected Alus – average of 100 iterations). Heatmaps further showed that a same set of first-Alus highly methylated in the long-lived donors contributed to the cycle of demethylation from newborn to middle-aged, followed by recovery in late life (boxed region, Figure 5D, and example Figure 5E). These observations suggested that the boundary function of first-Alus is subject to regulation early in life, while possibly deregulated upon ageing. As transcription factor binding sites are also present upstream of first-Alus, we speculate that differential methylation of regions upstream of genes may contribute to regulating access of these binding sites (example Figure 5E, bottom track).

We finally used this data set to gain information on the dynamic of the H3K4me1 signal. The initial study had defined marker genes for newborn, middle-aged, and long-lived donors (41). This allowed us to examined the distribution of the H3K4me1 signal in adult E034 T cells at genes active in middle-aged adults, or having been active earlier in life, or not yet at their peak activity. Predictably, the H3K4me1 signal was strongest at genes highly expressed in the middle-aged, but concentrated in a peak on the first-Alu at genes displaying maximum expression earlier in life (Figure 5F, dark and light blue profiles). Inversely, we did not observe any peak at the few late-life marker-genes (Figure 5F, yellow profile).

Together, these observations suggest that first-Alus form a dynamic boundary for DNA methylation, while they also function as memory media, recording earlier phases of transcriptional activity, via positioning of H3K4me1 histone modifications.

## Discussion

In the human genome, Alu elements are most abundant in the euchromatic interLADs, rich in genes. This enrichment contrasts with the scarcity of these repeats in the regulatory elements (REs) controlling these genes. Intuitively, this counterselection of Alu elements in REs is explained by the need to preserve the integrity of transcription factor binding sites and other DNA motifs involved in the regulation of transcription initiation. Alternatively, locating Alu repeats at the periphery of REs may participate in fulfilling unexplored requirements of the genes. Here, we bring several observations in favor of this second, not exclusive possibility.

Firstly, in T cells, we noted a very clear match between the profiles of Alu distribution and DNA methylation upstream of genes, with the “methylation landscape” transiting from “valleys” to “peaks” at the first Alu encountered by the RNA PolII after the TSS (referred to as the first-Alu, see models Figure 6A and 6B, black profiles). This upstream DNA methylation matching the local Alu density was independent of the DNA methylation at core promoters, detected only at transcriptionally inactive genes. The upstream DNA methylation may be involved in avoiding spurious transcription initiation outside core promoters, alike what is observed inside genes (7). In this context, we speculate that the drifting of the DNA methylation limit from the first-Alu to more upstream regions between newborn and middle-aged individuals may allow transcription factors to gain access to binding sites located upstream the first-Alus when the immune system becomes mature (see model Figure 6C). In the long-lived donors, we observed a genome-wide increase in DNA methylation specifically affecting Alu elements and occurring at first-Alus more abundantly than expected by chance. There are two possible interpretations of this. Firstly, the regain in methylation may interfere with transcription factor binding in aged individuals and compromise gene activity. But most studies on aging report a loss of methylation, in the context of an erosion of chromatin compartments (48). Therefore, a second possibility is that the long-lived, almost centenarian donors having contributed to the study may have aged particularly well. If this is the case, the increased methylation at Alus may possibly have a protective effect against spurious gene activity.

**Figure 6:**
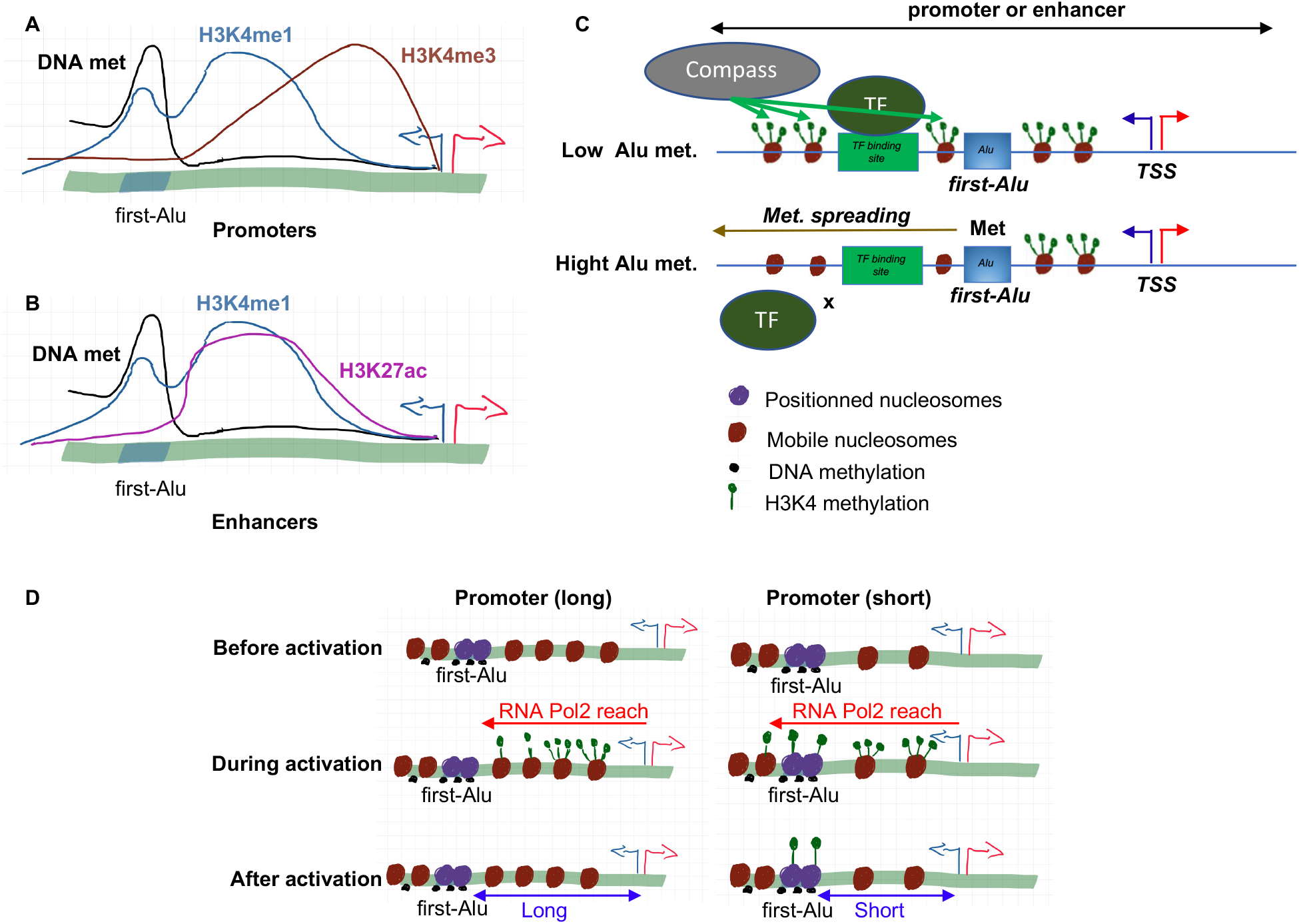
Models. (A-B) Model part 1: At promoters, the first-Alu sets the limit between the upstream DNA methylation landscape and the promoter-borne H3K4me3 signal. Unlike the H3K4me3 signal, the H3K4me1 crosses the first-Alu; this mark also accumulates at the first-Alu. At enhancers, DNA methylation and H3K4me1 distribute as described for promoters. H3K27 acetylation accumulates mostly in the interval between TSS and first Alu, but eventually spreads beyond the Alu boundary. (C) Model part 2: when first-Alu methylation is low (*i*.*e*. affecting only a small fraction of the cell population), transcription factor binding sites upstream of the first-Alu are located in an area carrying H3K4me3 marks and allowing for transcription factor binding and/or activity. When first-Alu methylation is high, the active promoter region stops at the first-Alu, and upstream transcription factor binding sites are out of commission. (D) Model part 3: nucleosomes are poorly positioned on the promoter region, but well-positioned at the first-Alu (line 1). At promoters with long TSS-to-first-Alu intervals, the RNA PolII rarely reaches the nucleosomes positioned on the first-Alu and both H3K4me3 and H3K4me1 locates to poorly positioned nucleosomes. In contrast, when TSS-to-first-Alu intervals are short, the RNA PolII reaches the nucleosomes positioned on the first-Alu and favors their H3K4me1 modification. DNA methylation, by interfering with recruitment of the histone methyltransferase, then prevents transition from H3K4me1 to H3K4me3 at these nucleosomes and thereby spreading of the H3K4me3 mark (line 2). After the phase of activity, nucleosome replacement gradually erases most H3K4me1 marks. But on the first-Alu, where nucleosomes are stably positioned, the modification persist long after transcription has ceased. Due to the presence of these histone marks, previously active promoters with short TSS-to-first-Alu intervals are poised for later reactivation (line 3).

The first-Alus were also limits for the H3K4me3 signal (Figure 6A, brown profile). At most promoters, this mark of transcriptional activation did not reach all the way from the TSS to the first-Alu, but when this encounter occurred, the H3K4me3 signal abruptly declined at this boundary. A similar boundary effect was observed at enhancers. In contrast, the H3K4me1 signals readily traversed the first-Alu boundary (Figure 6A and 6B, blue profiles), while the H3K27ac mark displayed an intermediate distribution, traversing the first-Alu but maintaining a clear enrichment on the inner side of enhancer CIAs (Figure 6B, purple profile). This distribution of the H3K4me3, H3K4me1, and H3K27ac marks, at large explains the systematic exclusion of Alu repeats from promoters annotated by the NIH Roadmap Epigenomics Mapping Consortium, as these are predicted by the chromHMM algorithm for which H3K4me3 levels have a high weight in the attribution of a promoter-type “chromatin state” (49). In contrast, the compatibility of Alu sequences with H3K4me1 and H3K27ac, mostly attributed to enhancers, would allow Alu elements to occasionally be annotated as enhancers.

In T cells and to a lesser extend in other cell types, the H3K4me1 signal underwent a peak of amplitude when crossing a first-Alu. This positioning of H3K4me1 at Alu elements close to promoters is a previously described phenomenon (13). When considered in the light of the several reports on nucleosome positioning at Alu repeats, it seems likely that this peak of signal corresponds to a phasing of H3K4me1-modified nucleosomes at that position. As Alu elements have an average length of 300 nucleotides, first-Alus would in average accommodate two nucleosomes, each with a footprint of 146 nucleotides (model Figure 6D, purple nucleosomes). Importantly, we further showed that intensity of the H3K4me1 peak gradually decreased as the TSS-to-first-Alu distance increased. In parallel, we showed that increased TSS-to-first-Alu distances correlated with decreased transcription of the first-Alu. Together, these observations suggest that transcription favors the mono-methylation of the nucleosomes, including those positioned on the first-Alus when they are within reach of the RNA PolII (Figure 6D, lines 1 and 2). A link between RNA PolII transcription and H3K4me1 deposition is fully compatible with earlier observations showing that the polymerase precedes the H3K4me1 at REs and that inhibiting RNA PolII elongations reduces levels of this modification at enhancers (50).

Examination of multiple tissues revealed that the average TSS-to-first-Alu distance at active promoters varied extensively from one tissue to the other, being highest in hematopoietic tissues and lowest in many immature cell types. This coincided well with the clear first-Alu-centered H3K4me1 peak observed in T cells, contrasting with a complete absence of peak in many tissues with larger average TSS-to-first-Alu distances. However, there was no systematic correlation, and signs of first-Alu-centered H3K4me1 peaks were observed in multiple non-hematopoietic tissues, possibly as a consequence of high transcriptional activity. Interestingly, the H3K4me1 mark was previously shown to accumulate at promoters as a consequence of temporary transcriptional activity (46), and the H3K4me1 mark was reported to be an indicator of a chromatin state poised for transcription (51). In that context, we find that marker genes of early life examined in mid-aged donors display stronger first-Alu-positioned H3K4me1 peaks than do middle-age marker genes, while the few marker genes for late life did not display this peak at all. This would be compatible with a local mono-methylation of the nucleosomes during periods of maximum transcription, and then a preservation of these marks at the first-Alu later in life, as the consequence of the nucleosome-positioning properties of Alu DNA sequences (Model Figure 6D, second and third line). As short TSS-to-first-Alu distances were frequently observed at immune genes, this first-Alu-positioned H3K4me1 signal remnant in middle-aged donors may function as a memory mechanism and possibly allow rapid reactivation of these genes. It may therefore be a manifestation of the concept of ‘‘Trained immunity’’ referring to immune cells becoming adapted to a certain stimulus and then responding in a stronger manner upon a second exposure (52). The fact that first-Alu-positioning of the H3K4me1 signal seems tuned down in various stem cells, while exacerbated in T cells would further be consistent with pluripotency calling for as little memory as possible, while immunity greatly benefits from training.

The mechanism allowing for the boundary function of the first-Alu is still an open issue. Yet, we speculate that the two different properties congregating on the first-Alus, namely a richness in CpGs and the presence of nucleosome positioning DNA sequence patterns, could be at the source of their boundary activity. Indeed, the primary depositors of global H3K4 trimethylation are the SET1A/B and MLL1/2 complexes. These two “COMPASS” complexes both harbor zinc finger-CXXC-domain proteins specifically binding unmethylated CpGs and participating in their targeting to promoters driven by CpG-islands (53). Examining CXXC1 associated with the SET1A/B complex yield a distribution consistent with a role for first-Alus in interrupting recruitment of this protein. DNA methylation at the first-Alu may therefore prevent recruitment of appropriate COMPASS methyltransferases and thereby interfere with the transition from H3K4me1 to H3K4me3 at the stably positioned nucleosomes. This would mechanically stop spreading of H3K4me3 beyond the first-Alu, and thereby create the boundary effect. In this scenario, persistent H3K4me1 at the first-Alu would be a secondary benefit of the Alu nucleosome positioning activity, serendipitously creating a memory effect.

Alu elements are not *per se* invaders of the human genome as they evolved from the 7SL gene, encoding an abundant cytoplasmic RNA participating in protein secretion (54). As such, they are legitimately involved in multiple functions for the benefit of the human genome (51). We here describe a new function for these repeated elements, participating in the definition of REs and being inflexion points for both DNA methylation and histone modifications indicative of transcriptional activity. We note however that relying on Alu elements as boundary and storage material may be the Achille’s heel of immune cell memory, as it is exposed to age-related decay in the form of modified DNA methylation at DNA repeats.

---

**Sup. Figure 1:**
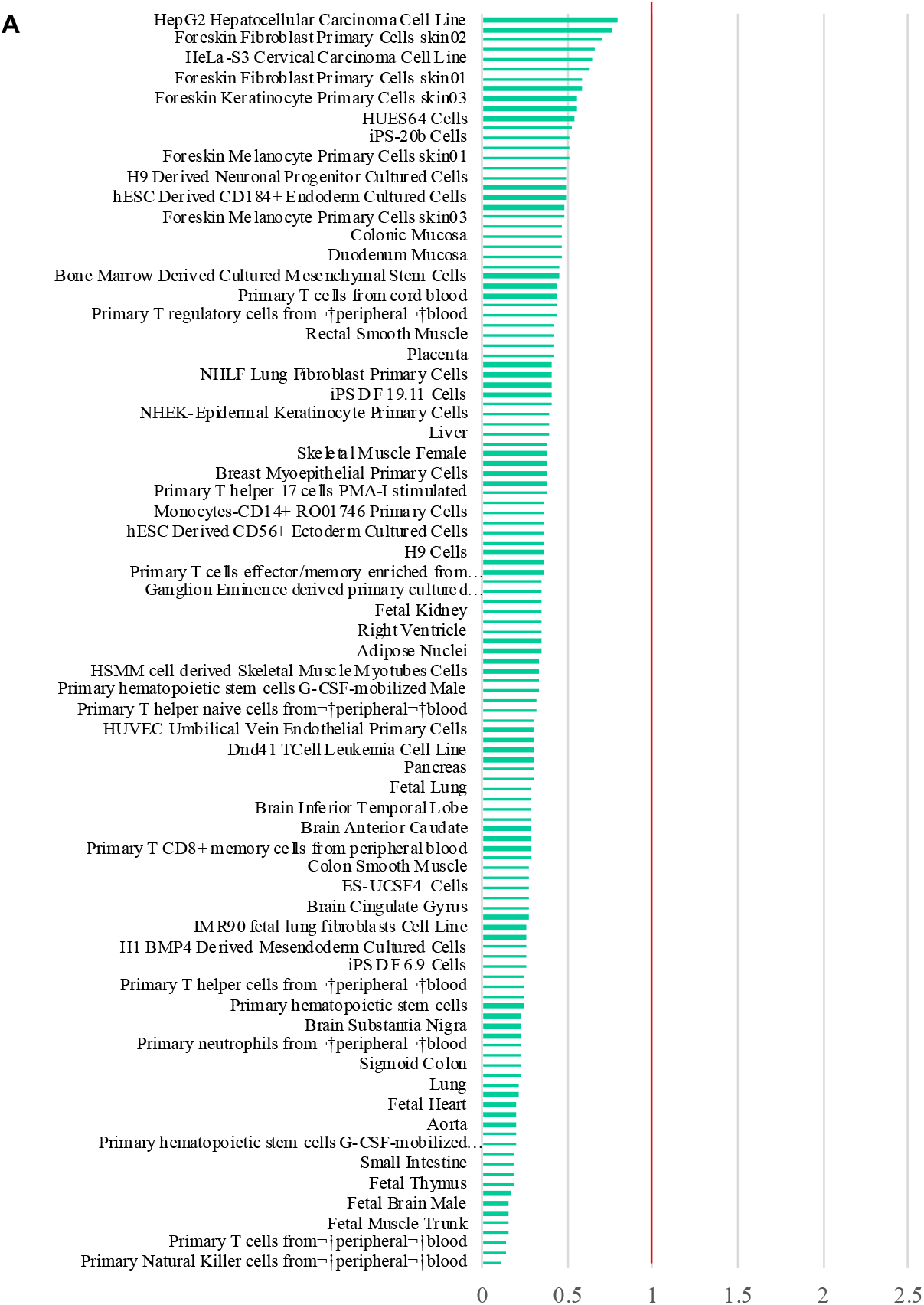
Tissue-specificity in the positioning of Alu elements relative to REs. (A) Comparison of regions annotated “Alu” in RepeatMasker with the regions annotated “1_TssA” in the 15 core marks model of the Epigenomic Roadmap consortium. For each tissue, the Jaccard index comparing enhancers to Alus is divided by the average Jaccard index (1000 iterations) obtained when comparing enhancers to randomly selected genomic locations (of the same sizes as the Alus). Only the tissues in the first and the last quartiles with the highest and the lowest scores are shown.

**Sup. Figure 2:**
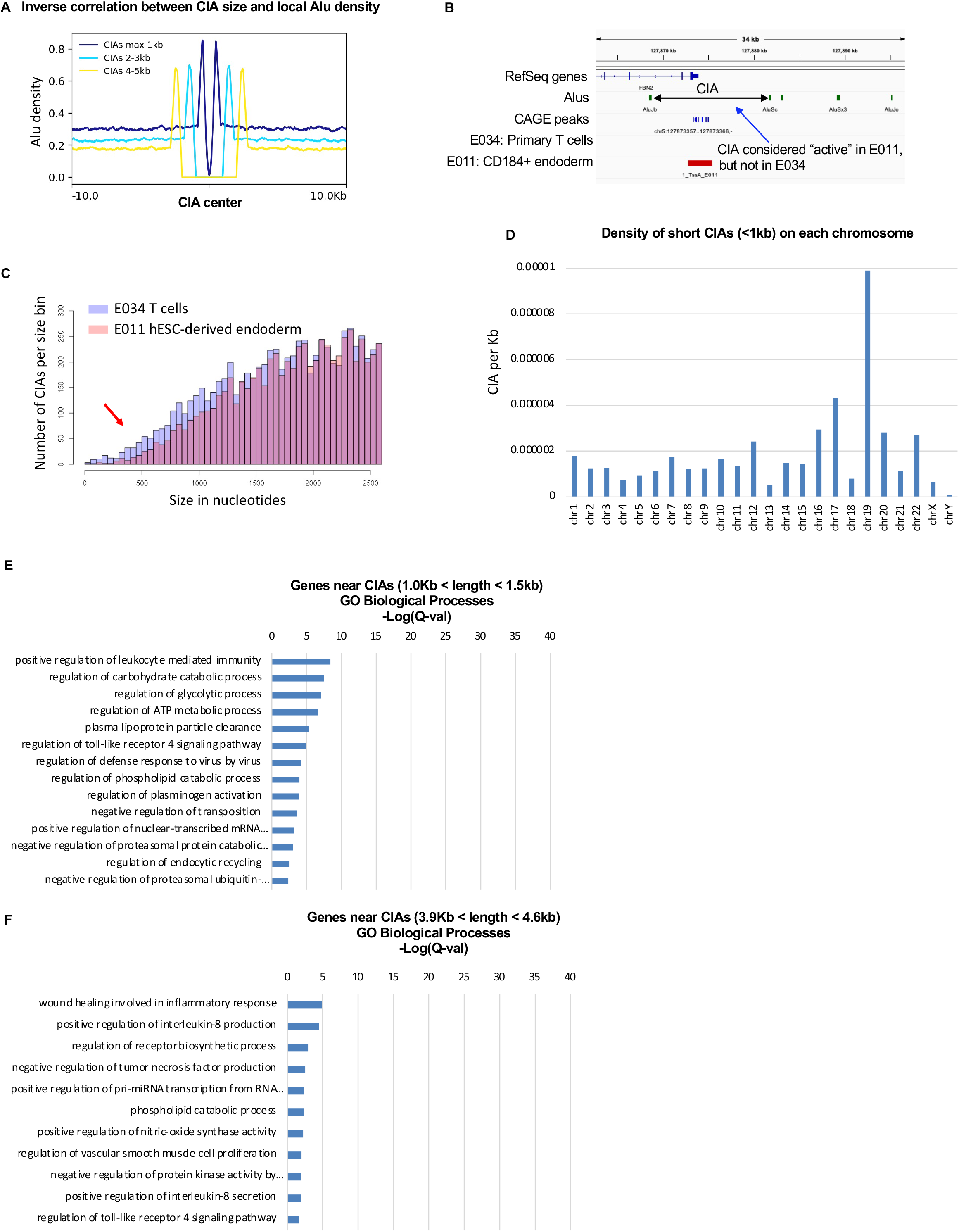
RE in regions of high Alu density are a specificity of immune genes. Average distribution Alu elements at CIAs of the indicated size (CIAs are regions located in-between two Alu elements and overlapping with a site of transcription initiation). The profiles are anchored on the center of the CIAs. The profiles illustrate that CIAs of increasing sizes translate into decreasing local Alu densities. (B) CIAs were considered “active” in a given tissue when they overlapped with an enhancer or a promoter annotated by the NIH Roadmap Epigenomics Mapping Consortium. In the given example, the CIA containing the TSS of the FBN2 gene overlapped with a region annotated as a promoter in E011 hESC-derived endoderm (red bar). It was therefore considered as active in that tissue. In E034 T cell, the region was not annotated, and therefore considered as inactive. (C) Bar-graph representing the CIA size distribution (in 50-nucleotide bins) in E034 T cells (blue bars) and E011 hESC-derived endoderm (red bars). Graph shows a bias toward small CIAs in E034 T cells (red arrow). (D) For each chromosome, <1kbCIA-density per chromosome was calculated by dividing the number of <1kbCIAs by the size of the chromosomes. (E) As a first control for the 4981 <1kbCIAs, we ranked all CIAs by size and selected the first 4981 CIAs with a size larger than 1kb (ranging from 1.0kb to 1.5 kb). Genes located in the neighborhood of these CIAs were identified using GREAT. The list of genes was then analyzed for enrichment in GO terms. Histogram shows the false discovery rate as -Log(Q value) as calculated by GREAT. (F) As a second control for the 4981 <1kbCIAs, we selected among the CIAs ranked by size, the 2490 CIAs smaller than the median (4.2kb) and the 2490 CIAs larger than the median (CIAs ranging from 3.9Kb to 4.6kb). Genes located in the neighborhood of these CIAs were identified using GREAT. The list of genes was then analyzed for enrichment in GO terms. Histogram shows the false discovery rate as -Log(Q value) as calculated by GREAT.

**Sup. Figure 3:**
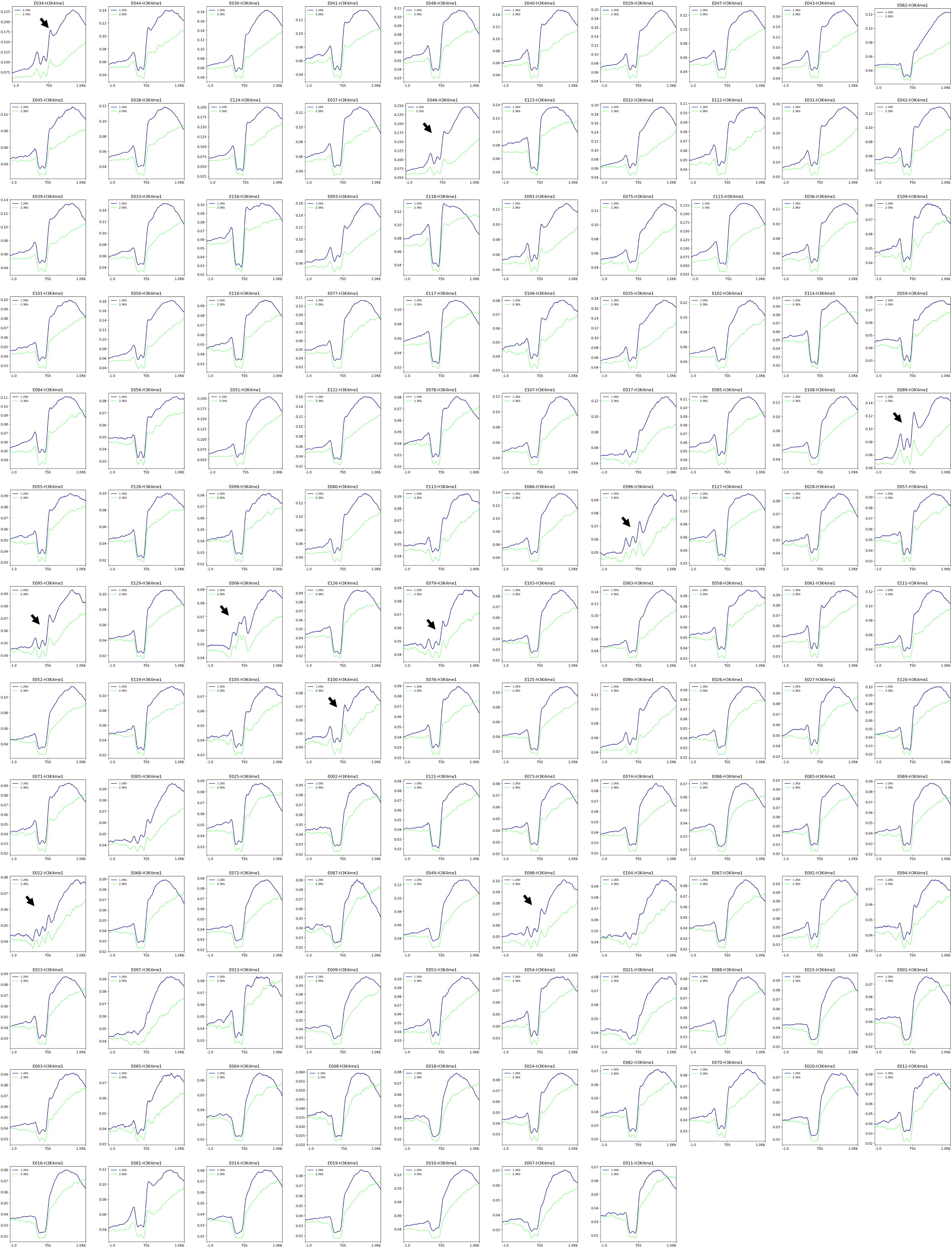
Peaks of H3K4me1 at first-Alus are detected in several tissues. For each of the 127 tissues annotated by the NIH Roadmap Epigenomics Mapping Consortium, profiles of H3K4me1 signal anchored on the first-Alus, for TSS-to-first-Alu intervals having a length of either 1 to 2kb, or 2 to 3kb. Arrows indicate unambiguous peaks of H3K4me1 signal at first-Alus. Tissue were labelled with the number given by the NIH Roadmap Epigenomics Mapping Consortium (see supplemental Table 1). “TSS” indicates the 3’-end of the first-Alu.

**Sup. Figure 4:**
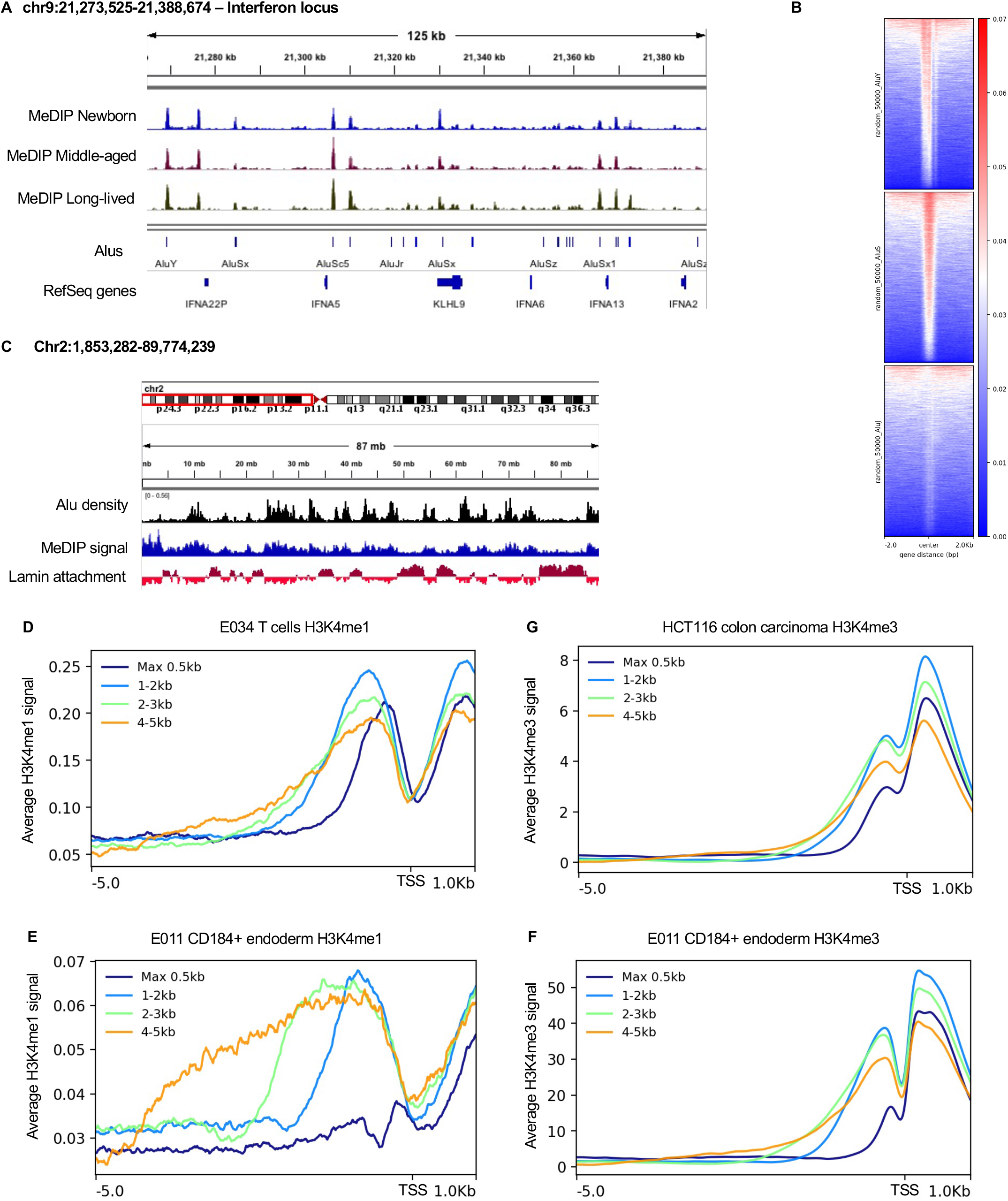
First-Alus are inflecFon point for epigeneFc marks at regulatory elements. (A)Screenshot from IGV – Example of the interferon locus. Paired-end MeDIP sequencing data (N=3 for each age) was aligned on the human genome with a requirement for unambiguous mapping of at least one of the ends to allow estimating DNA methylation in the immediate vicinity of repeated DNA elements, while eliminating all multi-mapping reads. (B) Heatmaps of the MeDIP signal at series of 50000 randomly selected Alu elements either having the most recently colonized the human genome (AluY), older Alu elements (AluS), or most ancient Alu elements (AluJ). (C) Screenshot from IGV – High Alu density and high MeDIP signal in regions not involved in lamin ahachment (interLADs - represent by negative red signal at the bohom track). Example from the short arm of chromosome 2. (D-G) For the indicated epigenetic marks, and the indicated cell types, profiles anchored on the TSS, for TSS-to-first-Alu intervals having a length of either less than 0.5kb, 1 to 2kb, 2 to 3kb, or 4 to 5kb.

**Sup. Figure 5:**
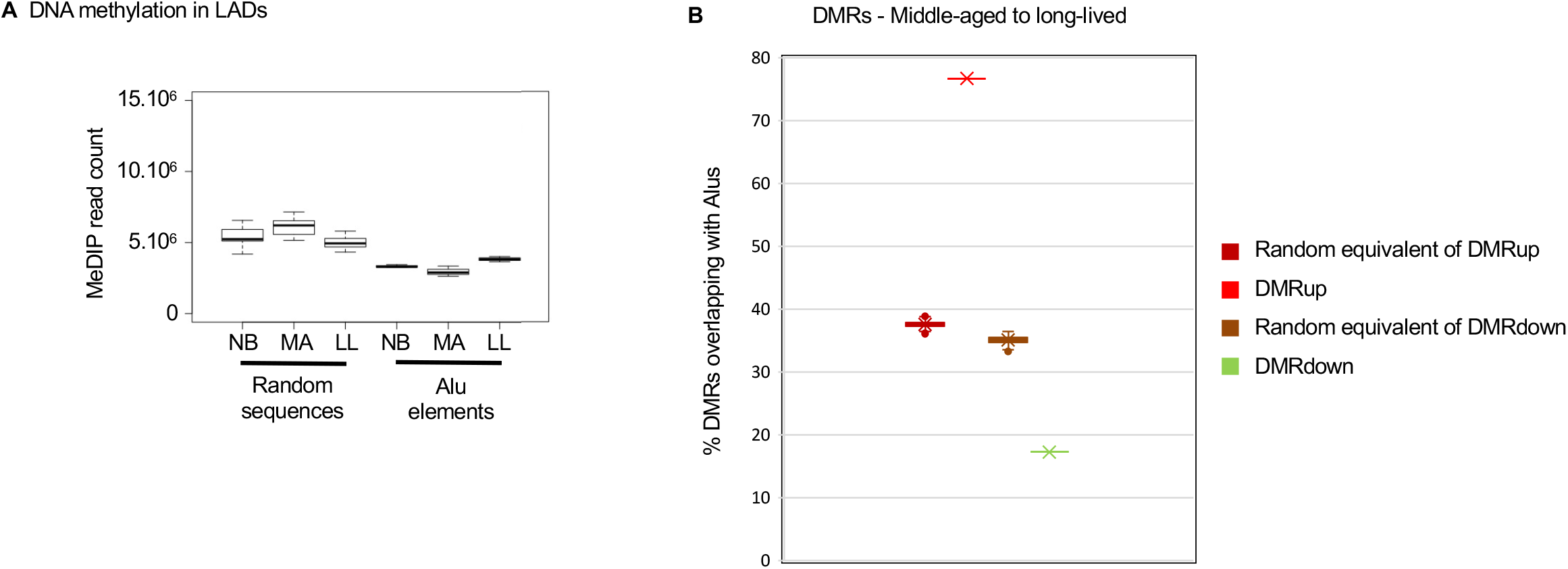
First-Alu DNA methylaFon and H3K4me1 peaking fluctuates with age. Methylation inside Alu elements or randomly selected regions in LADs at three different ages: counting of MeDIP reads either mapping to Alu elements present within LADs or mapping to an identical set of non-Alu LAD regions in the indicated age-groups. (B) Fraction (in %) of the middle-aged to the long-lived DMRs overlapping with Alu elements. Overlap expected by chance is also indicated (average of 10 iterations).

